# Comparative anatomy of leaf petioles in temperate trees and shrubs

**DOI:** 10.1101/2021.04.01.438018

**Authors:** Arinawa Liz Filartiga, Adam Klimeš, Jan Altman, Michael Peter Nobis, Alan Crivellaro, Fritz Schweingruber, Jiří Doležal

## Abstract

Petioles are important plant organs connecting stems with leaf blades and affecting light-harvesting leaf ability as well as transport of water, nutrient and biochemical signals. Despite petiole’s high diversity in size, shape and anatomical settings, little information is available about their structural adaptations across evolutionary lineages and environmental conditions. To fill our knowledge gap, we investigated the variation of petiole morphology and anatomy in 95 European woody plant species using phylogenetic comparative models. Two major axes of variation were related to leaf area (from large and soft to small and tough leaves), and plant size (from cold-adapted shrubs to warm-adapted tall trees). Larger and softer leaves are found in taller trees of more productive habitats. Their petioles are longer, with a circular outline, thin cuticles without trichomes, and are anatomically characterised by the predominance of sclerenchyma, larger vessels, interfascicular areas with fibers, indistinct phloem rays, and the occurrence of prismatic crystals and druses. In contrast, smaller and tougher leaves are found in shorter trees and shrubs of colder or drier habitats. Their petioles are characterized by teret outline, thick cuticle, simple and non-glandular trichomes, epidermal cells smaller than cortex cells, phloem composed of small cells and radially arranged vessels, fiberless xylem, lamellar collenchyma, acicular crystals and secretory elements. Individual anatomical traits were linked to different internal and external drivers. The petiole length and vessel conduit size increase, while cuticle thickness decreases, with increasing leaf blade area. Epidermis cell walls are thicker in leaves with higher specific leaf area. Collenchyma becomes absent with increasing temperature, epidermis cell size increases with plant height and temperature, and petiole outline becomes polygonal with increasing precipitation. We conclude that species temperature and precipitation optima, plant height, leaf area and thickness exerted a significant control on petiole anatomical and morphological structures not confounded by phylogenetic inertia. Unrelated species with different evolutionary histories but similar thermal and hydrological requirements have converged to similar petiole anatomical structures. Our findings contribute to improving current knowledge about the functional morphoanatomy of the petiole as the key organ that plays a crucial role in the hydraulic pathways in plants.

## 1. Introduction

Petioles are one of the most efficient structures in plants and represent an essential connection between the stem and the plant’s photosynthetic machinery, the leaf blade (Faisal et al. 2010). Petiole’s main function is to provide mechanical support to self-hold and adjust leaf position towards the sun, improving leaf ability of harvesting light. Besides, they also have a key role in the hydraulic pathway throughout the plant, transporting water, nutrients and biochemical signals to the leaves, and photosynthates and other products towards the shoot (Niinemets and Fleck 2002). Commonly green, and thus photosynthetically active, petioles may be stiff or flexible, long or short, but are usually wider at their base. In some woody taxa, they also can activate an abscission meristem that ensures proper leaf shedding at senescence. Similar to the leaf blade size variation (Wright et al. 2017), the wide variation of petiole size, toughness and lifetime among plants must be considered when specific metabolic activities are evaluated. Vapour loss and hydraulic conductivity, for instance, change not only among species but also within the same taxa and are dependent on leaf position, sun exposure and vascular features (Sack et al. 2005; Poorter and Rozendaal 2008).

Although petioles and leaves are highly diverse in size, shape and hence anatomical settings, comparative studies linking petiole structures with leaf parameters or whole plant size across species and evolutionary lineages are rare. Most studies involve only one or few closely related species (Niklas 1991; Aasamaa and Sõber 2010; Brocious and Hacke 2016), which provide a narrow view of potential variation. Petiole anatomical studies are mainly descriptive focusing on the characterization of distinct taxa (Metcalfe and Chalk 1950) or solving specific taxonomic problems (Ganem et al. 2019; Palacios-Rios et al. 2019; Karaismailoğlu 2020). Nevertheless, anatomical traits are frequently combined with ecophysiological aspects (Nicklas 1991, 1992; Tadrist et al. 2014). Information about petiole biomechanic is commonly used to explain relationships between structure, such as the size of epidermal cells and cross-sectional geometry, with the size of leaf blade, petiole stiffness, bending capacity, or leaf angle (Niinemets and Fleck 2002; Faisal et al. 2010; Levionnois et al. 2020). Additionally, distinct xylem traits such as vessel number, diameter and length have been used to examine physiological mechanisms of leaf hydraulic potential and vulnerability to xylem cavitation in petioles (Hacke and Sauter 1996; Coomes et al. 2008; Aasamaa and Sõber 2008; Hochberg et al. 2014; Brocious and Hacke 2016; Gebauer et al. 2019).

Little information is available about the relative importance of internal (leaf and plant size) and external climatic (temperature, precipitation) drivers for interspecific variation in petiole anatomical and morphological structures. Only a few studies provide an ecological interpretation that combines petioles morphoanatomy of one or a few species with abiotic conditions such as desiccation, light availability and wind gradient (Nicklas 1991; Hacke and Sauter 1996; Niinemets and Fleck 2002; Abrantes et al. 2013; Klepsch et al. 2016; Louf et al. 2018). Likewise, it is well-known that large-statured and large-leaved plants predominate in wet, warm and productive environments, while short-statured and small-leaved plants occur in cold or arid conditions such as high latitudes and elevations (Wright et al. 2017), but little is known about variation in petiole anatomy across these environmental and plant size gradients. Concerning vessel diameter variation, for instance, two explanations seem to complement each other, based on studies of roots, stems and branches (e.g., Zimmermann & Potter 1982; Alder et al. 1996; Hacke & Sauter 1996; Martinez-Vilalta et al. 2002). One posits that the variation in vessel diameter reflects environmental constraints operating at the whole-plant level (Hacke et al. 2016), while the other relates increasing vessel diameter along the plant stem to accomplish sap conduction requirements (Olson et al. 2013). However, to our knowledge, there are no studies until now that would combine petiole traits with whole-plant size, leaf traits and environmental factors in a phylogenetic context.

To fill this knowledge gap on petiole morphoanatomical trait variation and to better understand internal and external drivers and constraints in an evolutionary context, we investigated the variation of petiole morphology and anatomy among a census of European woody species growing in contrasting environmental conditions using phylogenetic comparative models (Adams et al. 2014). In particular, we studied how petiole anatomical features differed according to whole-plant size, leaf traits, thermal and hydrological conditions, and taxonomical origin for a wider group of temperate trees and shrubs. The study is based on 95 woody species and 19 morphoanatomical features, such as cross-sectional geometry, epidermis traits, conductive, mechanical and storage tissue structures, many of which serve as a proxy for ecophysiological adaptations which are more difficult to study for a large number of species. Given that the leaves are the main photosynthetic apparatus of plants and that the petioles have a major role to support them mechanically and physiologically, we assumed that the structure of the petiole will be more affected by leaf characteristics than by abiotic factors. We also expected that the supporting tissues (collenchyma and sclerenchyma) would be more developed in the larger leaves to optimize the ability to retain leaves. Besides, because leaf size variations are directly reflected in photosynthesis and water transport supply, we expected the vessel diameter to increase with the leaf blade area.

## 2. Materials and methods

### 2.1. Plant species

Our analysis was based on 95 woody species occurring in temperate and Mediterranean regions of Europe. The species belong to 72 genera and 35 families, with Rosaceae (14 taxa) as the most species-rich family, followed by Betulaceae (8), Salicaceae (7), Fabaceae (6), Fagaceae (6), Oleaceae (6), Caprifoliaceae (5), Ericaceae (5), Anacardiaceae (4), and Cornaceae (3). *Prunus* is the most represented genera (5), followed by *Alnus* (4), *Quercus* (4), *Salix* (4), *Populus* (4) and *Acer* (3). Most of the studied taxa are small trees (29), followed by large trees (25), large shrubs (19) and small shrubs (17), and woody lianas (5). Figure 1 shows examples of few species that we analyzed in this work.

**Figure 1.**
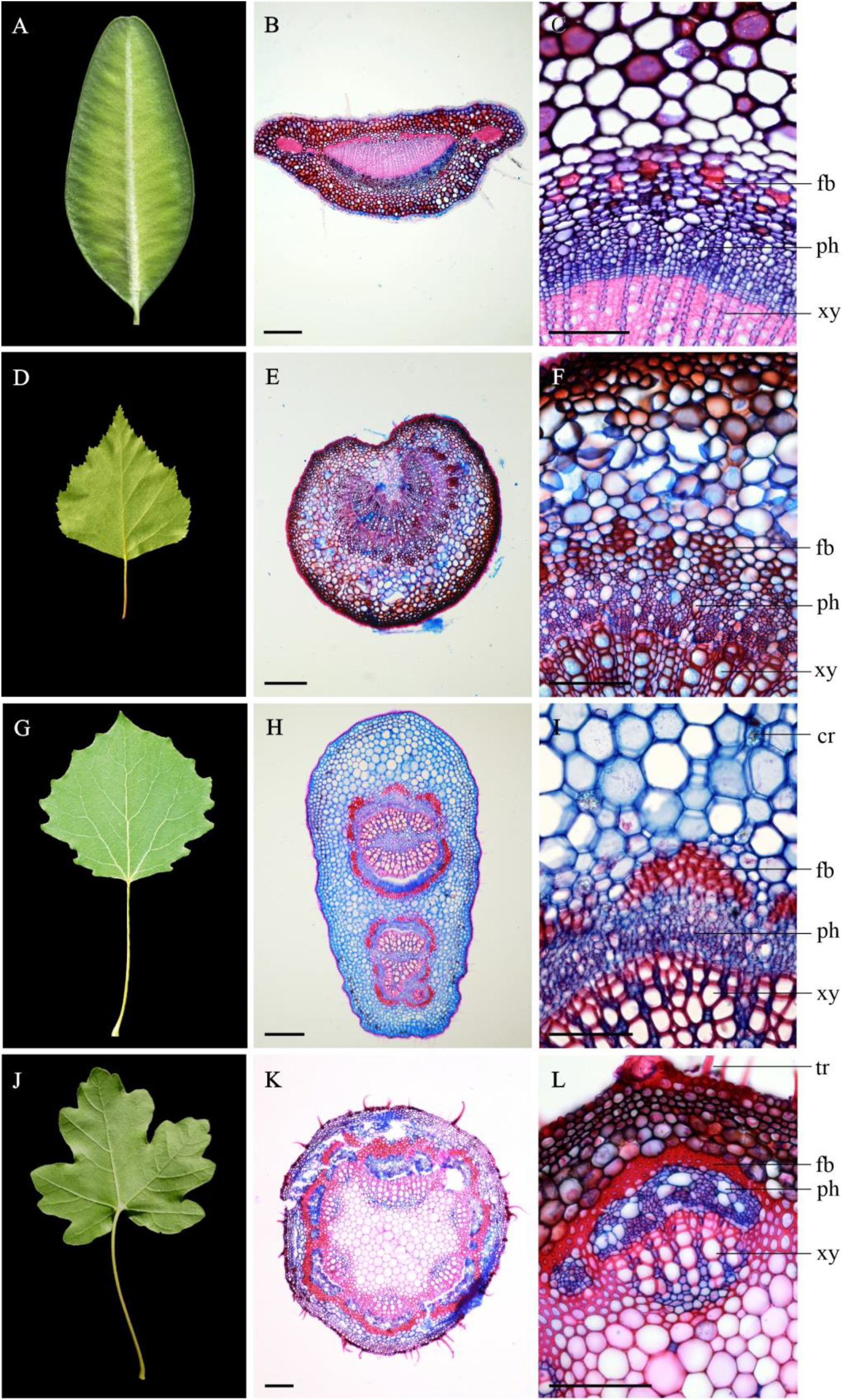
*Buxus sempervirens* (A-C), *Betula pendula* (D-F), *Populus tremula* (G-I), *Acer campestre* (J-L). Petiole length is classified in <2-5 cm (A), 5-10 cm (D, G) and >10 cm (J). Cross-sections show petiole geometry: horizontally flattened (B), with an indentation (E), vertically flattened (H) and circular (K); and point out trichomes (tr), fiber band (fb), phloem (ph), xylem (xy) and crystals (cr). Scale bars: 200 µm (B, E, H, K) and 100 µm (C, F, I, L). DNA (ITS). Varying mutational rates between loci maximize the ability to discern major lineages together with species-level phylogenies; this threesome of markers was therefore suggested as a standard for phylogenetic analyses and barcoding of angiosperms in the past (Li et al. 2017).

### 2.2. Morphology

For each species, we randomly selected and sampled 3-5 fully expanded and uninjured leaves. For most of the species, we obtained data of plant height, leaf dry matter content (LDMC), specific leaf area (SLA) from LEDA database (Kleyer et al. 2008). We acquired leaf area (LA) values from BIEN database (Enquist et al. 2016), TRY database (Kattge et al. 2020) and, in the case of *Cercis siliquastrum*, from the literature (Hatamian et al. 2019). For compound leaves, values for the whole leaves were used. We measured LA and LDMC for six species (i.e., *Alnus cordata, Magnolia soulangeana, Pyrus communis, Rhododendron ferrugineum, Ulnus glabra* and *Vitex agnus*) with no data available in LEDA. Leaf area measurements were done using the ImageJ software (Dolezal et al. 2019). Mean values per species were used in further analyses.

### 2.3. Petiole anatomy

Petioles were fixated and stored in 70% ethanol. Cross-sections 20-30 µm thick were made on the middle region of the petiole using a sledge microtome. We discolored the sections with sodium hypochlorite, then double-stained them with a 1:1 aqueous Astrablue (0,5%) and Safranin (1%) blend, and finally mounted on a permanent slide with Canada Balsam (Gärtner and Schweingruber, 2013; Schweingruber et al. 2020). The slides were examined using an Olympus BX53 microscope, Olympus DP73 camera, and cellSense Entry 1.9 software.

We described a total of 19 anatomical features including cross-sectional geometry (outline), epidermis traits (cell size and wall width), cuticle thickness; the presence of trichomes, hypodermis, collenchyma, fiber band, crystals and secretory structures; the arrangement of vascular bundles and interfascicular region, phloem (size of parenchyma cells) and xylem (vessel arrangement and diameter, presence of fibers and distinct latewood; Figure 1). The complete characterization of the evaluated traits is presented in the Supplementary material (Figures S1-S51).

Quantitative anatomical measurements were also performed for each species using the software ImageJ. Cuticle thickness was obtained with an average of 20 measurements made along the periphery of the petiole. The evaluation of tissue predominance was focused on the vascular system (phloem and xylem) and fiber band (sclerenchyma); the total area of each tissue was measured. For the average vessel diameter (vessel size), we measured all the vessels present in several pre-defined representative squared areas with 100 µm on each side and average values were then calculated.

### 2.4. Phylogeny

The phylogenetic relationships between the species studied were constructed based on three molecular markers: matK, rbcL, and ITS. Combined, these three loci cover protein-coding, RNA-coding, and noncoding sequences, as well as both plastid (matK, rbcL) and nuclear

As a starting point, we acquired all relevant sequences from NCBI GenBank. All three combined matrices (one dataset per locus) were afterward aligned in MAFFT 6 (Katoh and Toh 2008), using the L-INS-i algorithm. Partial alignments were concatenated, manually adjusted in BioEdit (Hall 1999), and subdued to the automated1 algorithm in trimAll software (Capella-Gutiérrez et al. 2009) to exclude highly divergent and gap-rich regions. The best-fit model for phylogenetic inference was selected according to the Bayesian information criterion (Schwarz 1978) using the Baseml core of Kakusan4 package (Adachi and Hagesawa 1996; Tanabe 2011), resulting in the choice of the GTR model with rate variation across locations simulated by a discrete gamma distribution (Γ8), autocorrelated by the AdGamma rates prior, and unlinked for particular gene partitions. This model was afterward submitted to MrBayes ver. 3.1.2 (Ronquist and Huelsenbeck 2003) as the basis for MCMC analysis, encompassing two independent runs with four Metropolis-coupled MCMC chains of 107 generations sampled after every 1000th generation. In each run, one Markov chain was cold and three were incrementally heated by a parameter of 0.3. The first 25% of entries were discarded as burn-in to eliminate trees sampled before reaching apparent stationarity, and the rest was used to compute the majority-rule consensus. The resulting tree contained nodes supported only by substituted sequences and hence had to be edited: internal phylogeny of these groups was collapsed to polytomy and branch lengths were averaged.

### 2.5. Data analyses

The estimation of species’ climatic optima was obtained by calculating species affinities for temperature and precipitation using the CHELSA climate database (Karger et al. 2016) in the combination with species geographic occurrences extracted from the GBIF database (Global Biodiversity Information Facility, www.gbif.org). To evaluate the effect of species’ temperature and precipitation preferences and plant morphology (height, LA, SLA and growth form) on their petiole anatomy, we first performed ordination of petiole anatomical features, using nonmetric multidimensional scaling, to find major axes of variation in anatomical features and to quantify the variation explained by environment and morphology.

Secondly, we used the phylogenetic distance-based generalized least squares model (D-PGLS, Adams 2014b) to quantify how much variation in anatomical settings is explained by LA and other main predictors (plant height, SLA, precipitation, and temperature) after accounting for LA while controlling for phylogenetic inertia. Our 19 anatomical variables were used to calculate the distance matrix among all 91 species (4 species excluded due to missing values) using a simple matching coefficient (Legendre and Legendre 1998): Distance = 1 – (Number of agreements / Number of variables)

The distance matrix was used as the response; LA, SLA, height, annual precipitation, and annual mean temperature were used as predictors. Since the distribution of most predictors was skewed or they had outlier values, we used the natural logarithm of leaf area, square root of height and for precipitation and temperature, we used the rank of their values. The used implementation of the model does not allow simultaneous estimation of the strength of the phylogenetic signal. We estimate it and account for it to the appropriate degree by running the model with a phylogenetic tree transformed with the varying value of Pagel’s lambda (Pagel 1999). We used lambda values from 0 (no phylogenetic signal) to 1 (phylogenetic signal corresponding to Brownian motion) by 0.01 and selected the model with the highest explained variability. Correlations among predictors were low (strongest was between leaf area and height 0.41), thus explained variability was not highly dependent on the order of the predictors – we present explained variability for leaf area and each other predictor after accounting only for leaf area and after accounting for all the other predictors. We also explored the effect of each of our predictors on each anatomical trait separately using the same modelling approach.

We also estimated the phylogenetic signal in the anatomy of petioles alone using the distance matrix which we had constructed for the main model. We estimated multivariate generalization of Blomberg’s K (Blomberg et al. 2003; Adams 2014a) and then data was tested by permutation if it was different from the Brownian motion model of evolution.

The D-PGLS model was evaluated and the phylogenetic signal was estimated and tested using package *geomorph* (version 3.3.0; Adams et al. 2019), phylogenetic tree was scaled using package *geiger* (version 2.0.7; Harmon et al. 2008). All analyses were done in R (version 4.0.0; R Core Team, 2020).

## 3. Results

In analyses of individual petiole features, the predictors leaf area, plant height, temperature, and precipitation were all linked to different anatomical traits (Figure 2). For instance, variation in vessel size and vessel arrangement was mainly linked to LA, epidermis cell size to plant height, the petiole length to LA, the cuticle thickness to SLA, the presence of trichome to LA. The epidermis cell wall thickness was mainly related to SLA variation. The collenchyma was strongly related to temperature, tissue type proportion was linked to precipitation, phloem rays to temperature, petiole outline to plant height and precipitation.

**Figure 2.**
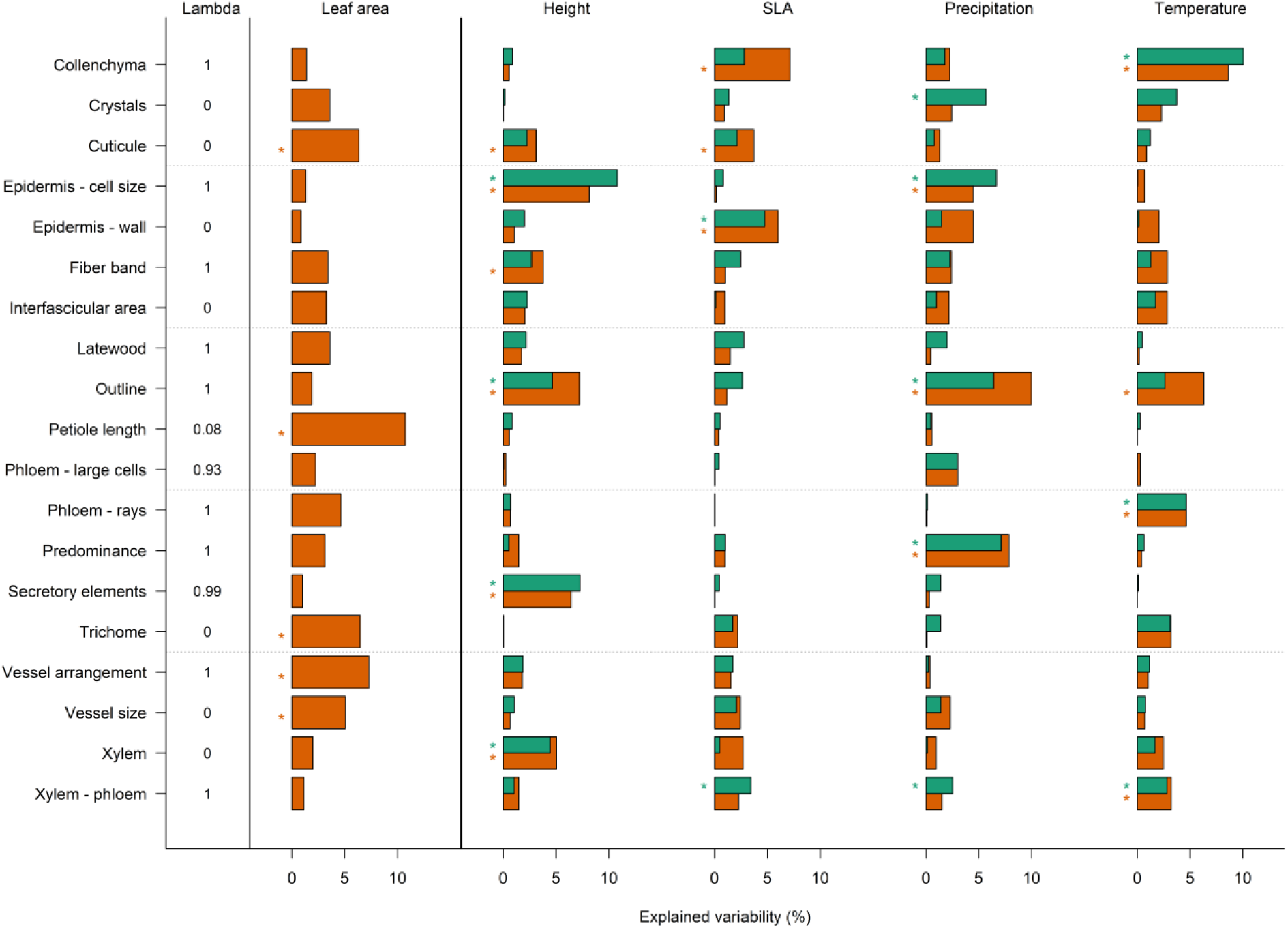
Explained variability in individual anatomical traits by LA, height, SLA, precipitation, and temperature. Brown bars show explained variability accounting only for the covariate LA; green bars denote explained variability after accounting for the other predictors (including LA). Stars show the cases with a p-value below 0.05. In the left column is the estimated strength of the phylogenetic signal in each model (Pagel’s lambda). LA = leaf area, SLA = specific leaf area.

Leaf area, height, SLA, precipitation, and temperature each affected petiole anatomy (p-values after accounting for other predictors: 0.028, 0.015, 0.035, 0.006, 0.008) and explained together 12.19% of the variability after controlling for phylogeny. The estimated phylogenetic signal of the main model was 1 (Pagel’s lambda) which corresponds to the Brownian motion model of evolution. The estimated strength of phylogenetic signal K_mult_ in anatomical traits only (without predictors) was 0.3983 which was significantly different from 1 (p-value 0.001). *Kmult* lower than 1 means the power of phylogenetic signal lower than expected under the Brownian motion model of evolution.

NMDS ordination (Figure 3) showed that the main interspecific differences in petiole anatomy along the first axis were associated with variation in leaf area (LA is positively correlated with SLA), from soft leaves with high SLA to tough leaves with low LA and high LDMC (SLA is negatively correlated with LDMC, a result not shown). The petiole anatomical changes along the second axis were associated with variation across temperature, precipitation, and plant height gradients, from cold-adapted shrubs (such as families Cornaceae, Grossulariaceae, and Aquifoliaceae) to warm-adapted tall trees (such as families Sapindaceae and Hippocastaneaceae) (Figure 3).

**Figure 3.**
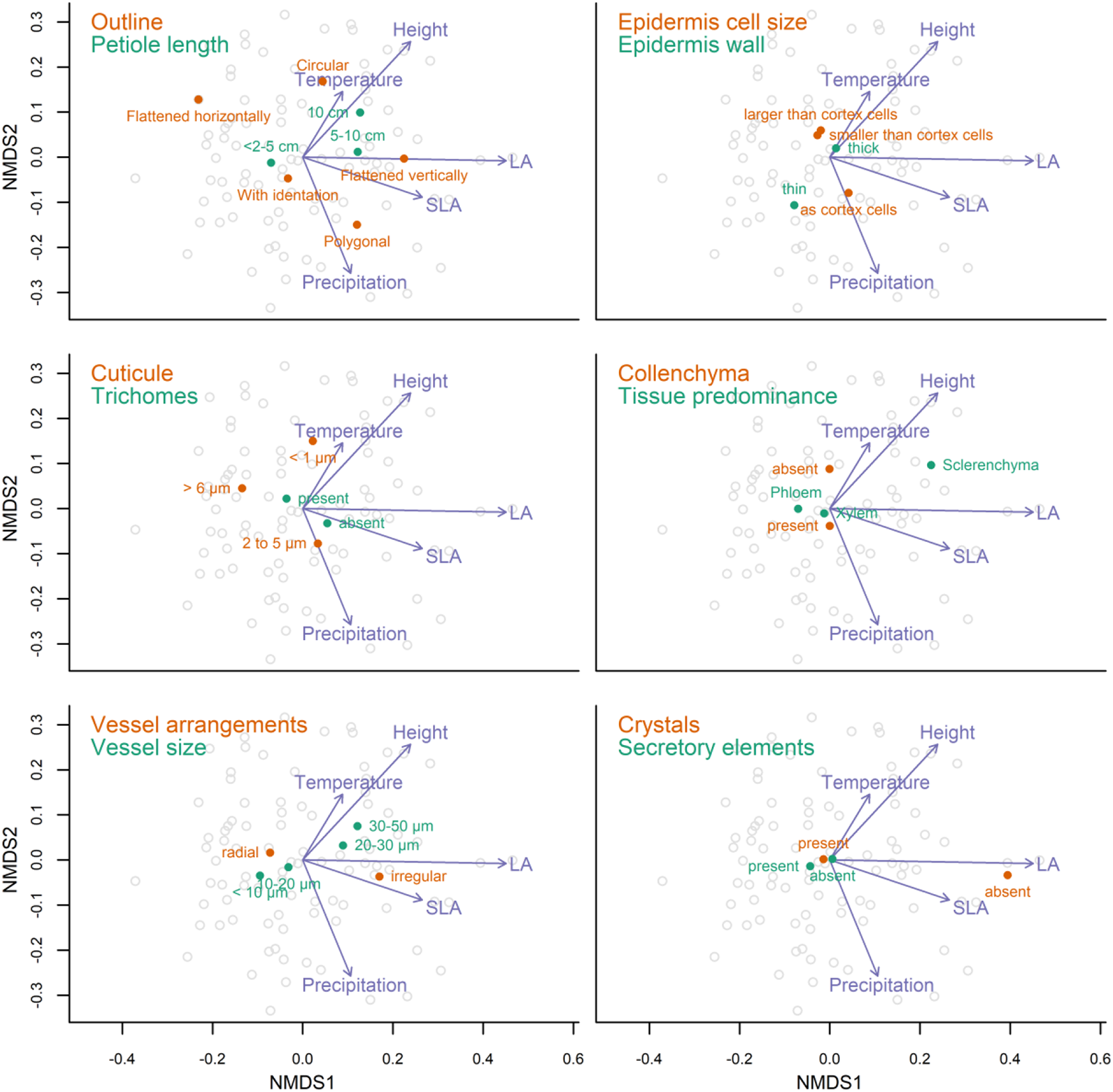
Ordination of species based on anatomical traits with a projection of predictors and selected anatomical traits. In all 6 panels, the same ordination is visualized only with projected centroids of levels of different selected traits. NMDS was done on distance matrix used in the main model (see methods), with 3 axes (only the first 2 are visualized) and with stress 0.175 (using package vegan (version 2.5-6; Oksanen et al. 2018)). NMDS = non-metric multidimensional scaling, LA = leaf area, SLA = specific leaf area.

Trees and shrubs with bigger and softer leaves (high LA, SLA, low LDMC) tended to have longer petioles (Figures 4 and 5) with epidermal cells of the same size as cortex cells, with thin cuticle without trichomes, irregularly arranged wide vessels, the predominance of sclerenchyma, phloem composed of large cells, and interfascicular areas with fibers. Petioles of large trees were usually longer (5-10 cm), with a circular external outline, thin cuticle < 1 µm, xylem fibers present, phloem rays indistinct, and prismatic crystals and druses (Figures 5 and 6). In contrast, trees with smaller leaves (low SLA and high LDMC) were rather of smaller stature from rather drier habitats with high precipitation seasonality and their petioles were characterized by thick cuticle > 6 µm, trichome simple and non-glandular, epidermal cells smaller than cortex cells, and phloem composed of small cells and radially arranged vessels (Figure 3). Moreover, petioles of smaller trees and shrubs tended to have an external form almost circular or with one flat part or with an indentation, cuticle 2 to 5 µm, fiberless xylem, fiber band absent, an interfascicular area without fibers, and lamellar collenchyma.

**Figure 4.**
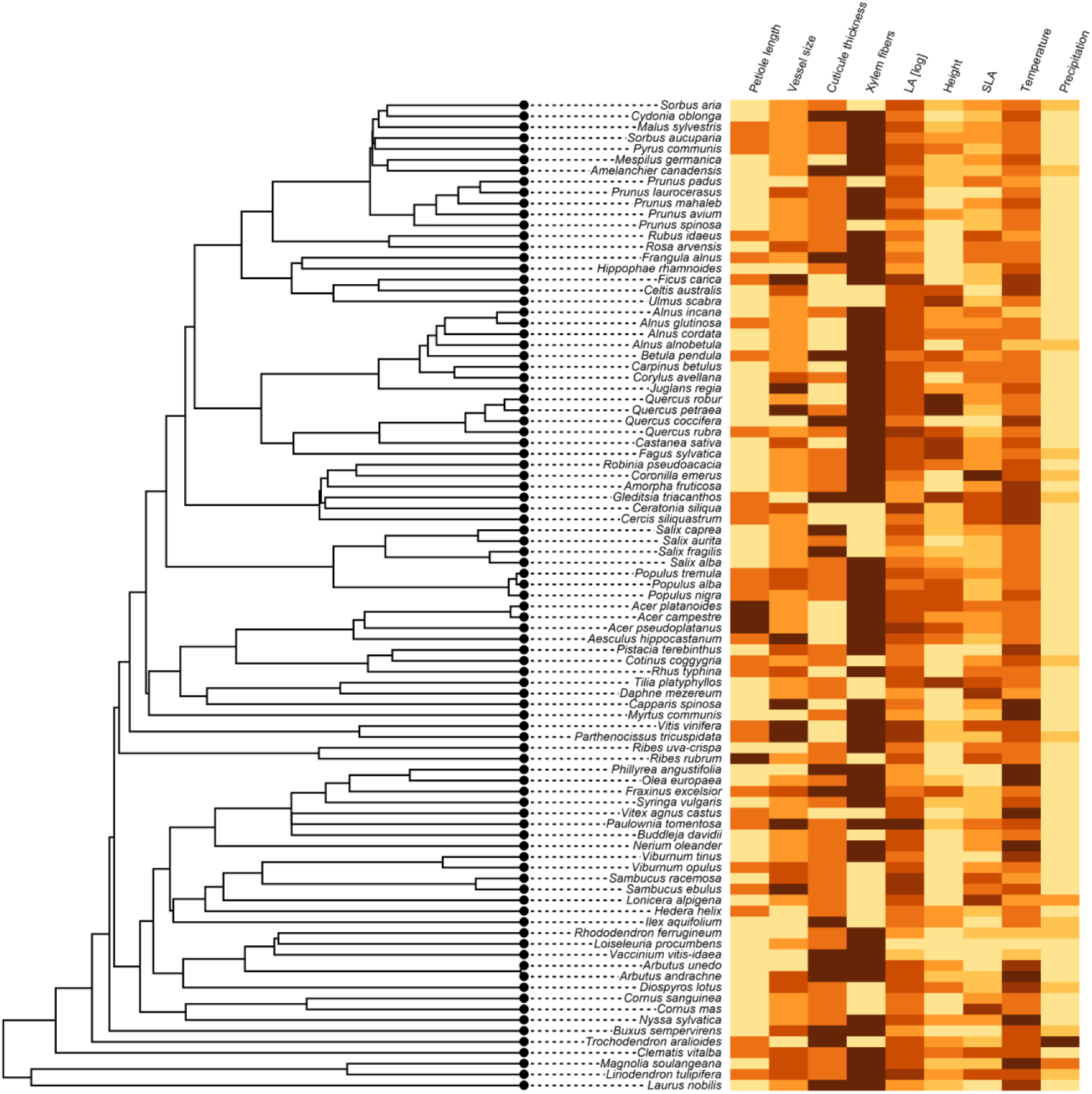
Phylogenetic tree with visualization of traits and environmental preferences of plants. The lowest values of each trait are in yellow and the highest values in brown.

**Figure 5.**
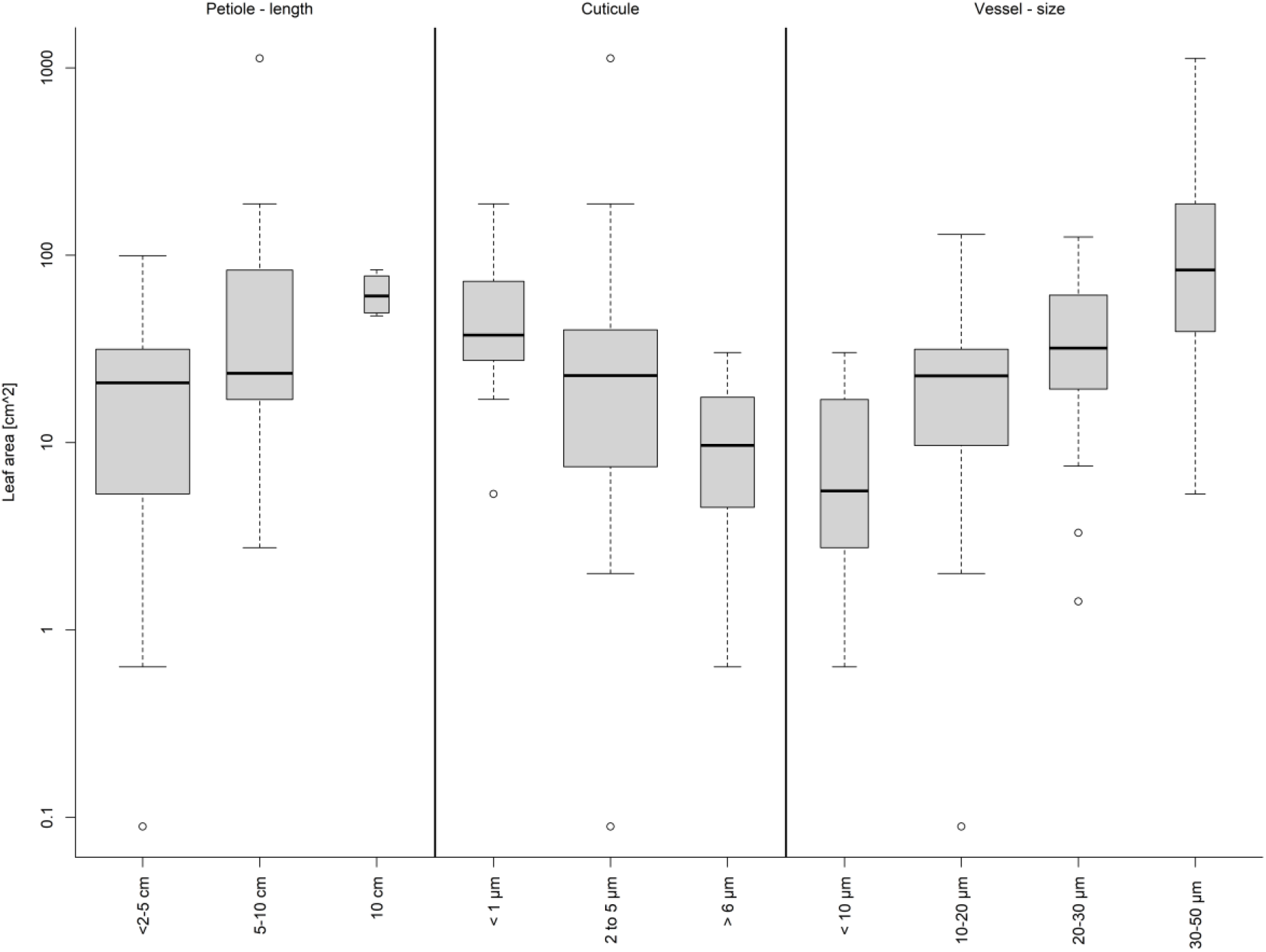
Relationship between leaf area and selected anatomical traits. The width of boxplots corresponds to the square root of the number of observations in a particular group.

**Figure 6.**
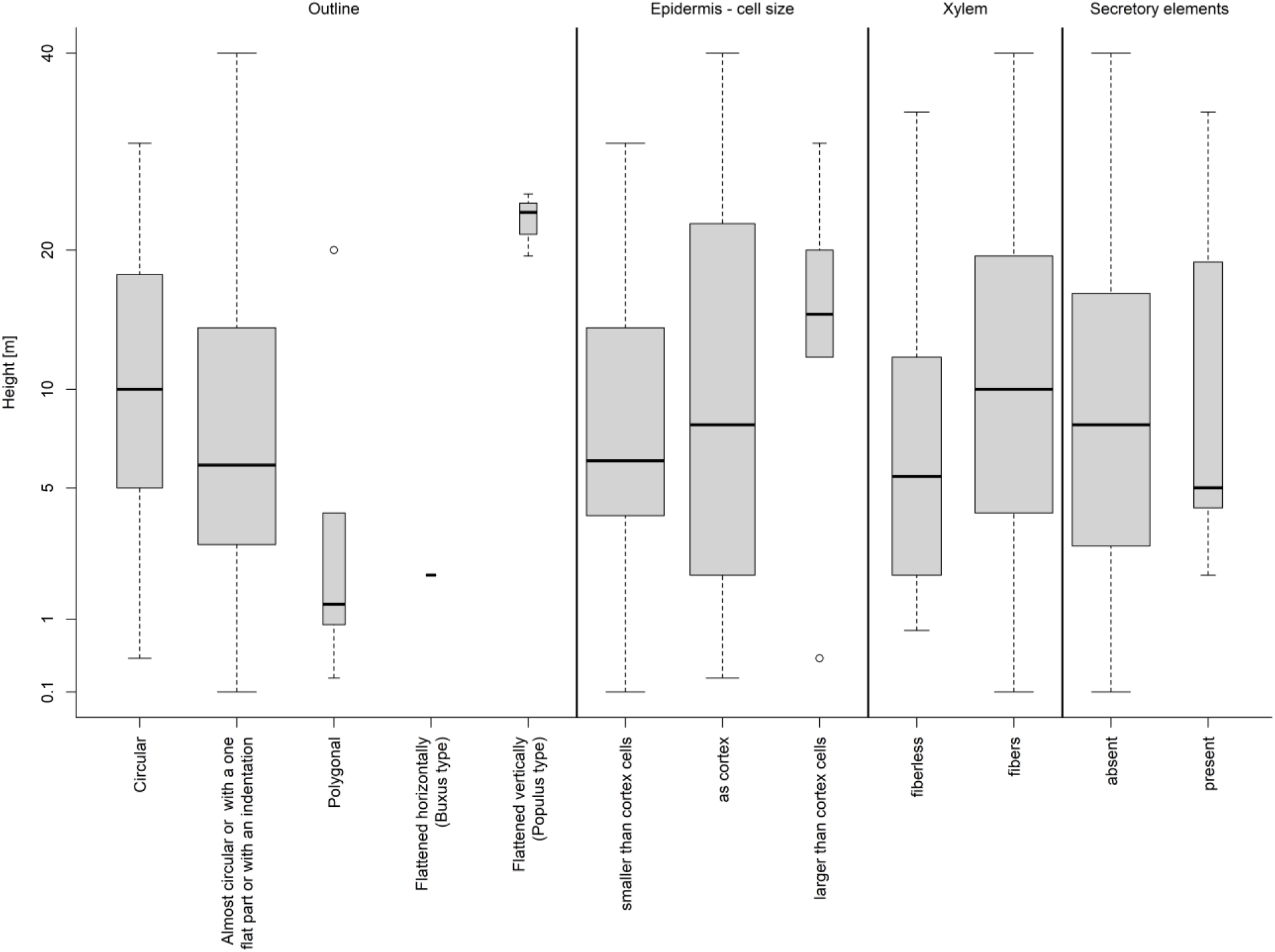
Relationship between height and selected anatomical traits. The width of boxplots corresponds to the square root of number of observations in a particular group.

### 3.1. Stabilizing elements: petiole outline, epidermis, collenchyma, and fiber bands

Plant height and precipitation accounted for the most variation in the petiole outline (Figure 3). The circular outline with flat section or indentation predominated among the studied species (68%), typically in shrubs and smaller trees from Cornaceae, Aquifoliaceae, Rosaceae, and Oleaceae. Circular petiole outline (22% of taxa) tended to predominate in tall trees from genera *Acer, Quercus, Tilia*, and *Carpinus* (Figure 6). The vertically flattened outline was found only in *Populus* spp. and horizontally flattened outline in *Buxus sempervirens*. Polygonal outline (5%) occurred in unrelated taxa such as *Clematis vitalba, Loiseleuria procumbens, Prunus laurocerasus, Ribes rubrum*, and *Sambucus ebulus* (Supplementary Material Figures S1-S5).

Unlike the petiole outline, the most variation in the petiole length was explained by the leaf area (Figures 2 and 5). Short petioles <2-5 cm occurred in 65% of species, followed by 5-10 cm long petioles (31%, Figure 6), while long petioles were found in only 4% of species including *Acer* spp. and *Ribes rubrum* (Figure 4).

Variation in petiole cuticle thickness was primarily related to LA (Figures 2 and 5), with shrubs and small trees of families Cornaceae, Aquifoliaceae, Rosaceae, and Oleaceae having smaller leaves with thick 2 to 5 µm cuticle, found in 49% of all species (Figure 4). The largest cuticle > 6 µm was characteristic of Mediterranean trees such as *Phillyrea angustifolia, Quercus coccifera* and *Laurus nobilis* (Figure 4). Thin cuticle < 1 µm predominated in large trees with bigger leaves such as *Acer spp, Carpinus betulus, Castanea sativa, Celtis australis, Ceratonia siliqua, Cercis siliquastrum*, and *Quercus robur* (Supplementary Material Figures S6-S14).

Plant height and precipitation accounted for the most variation in the petiole epidermal cells (Figures 2 and 6), which are in 56% of species smaller than cortex cells (Figure 5), usually in smaller trees or shrubs with small SLA such as *Salix, Prunus, Corylus, Vaccinium, Viburnum*, and *Syringa* (Figure 4). Tall trees of genera *Quercus, Populus, Ulmus*, and *Tilia* had petiole epidermal cells of the same size as cortex cells. Petiole epidermal cells were only in 6% of cases larger than cortex cells such as in *Aesculus hippocastanum* and *Prunus avium* (see also Supplementary Material Figures S6-S14). SLA accounted for the most variation in the epidermal cell-wall thickness (Figure 7). Most of the tree species with larger SLA had thick-walled epidermal cells (84%), while thin epidermal cell walls occurred among lianas (*Vitex, Hedera*) and shrubs *(Ilex aquifolium, Capparis spinosa, Viburnum* spp., *Glyzirhyza glabra*).

**Figure 7.**
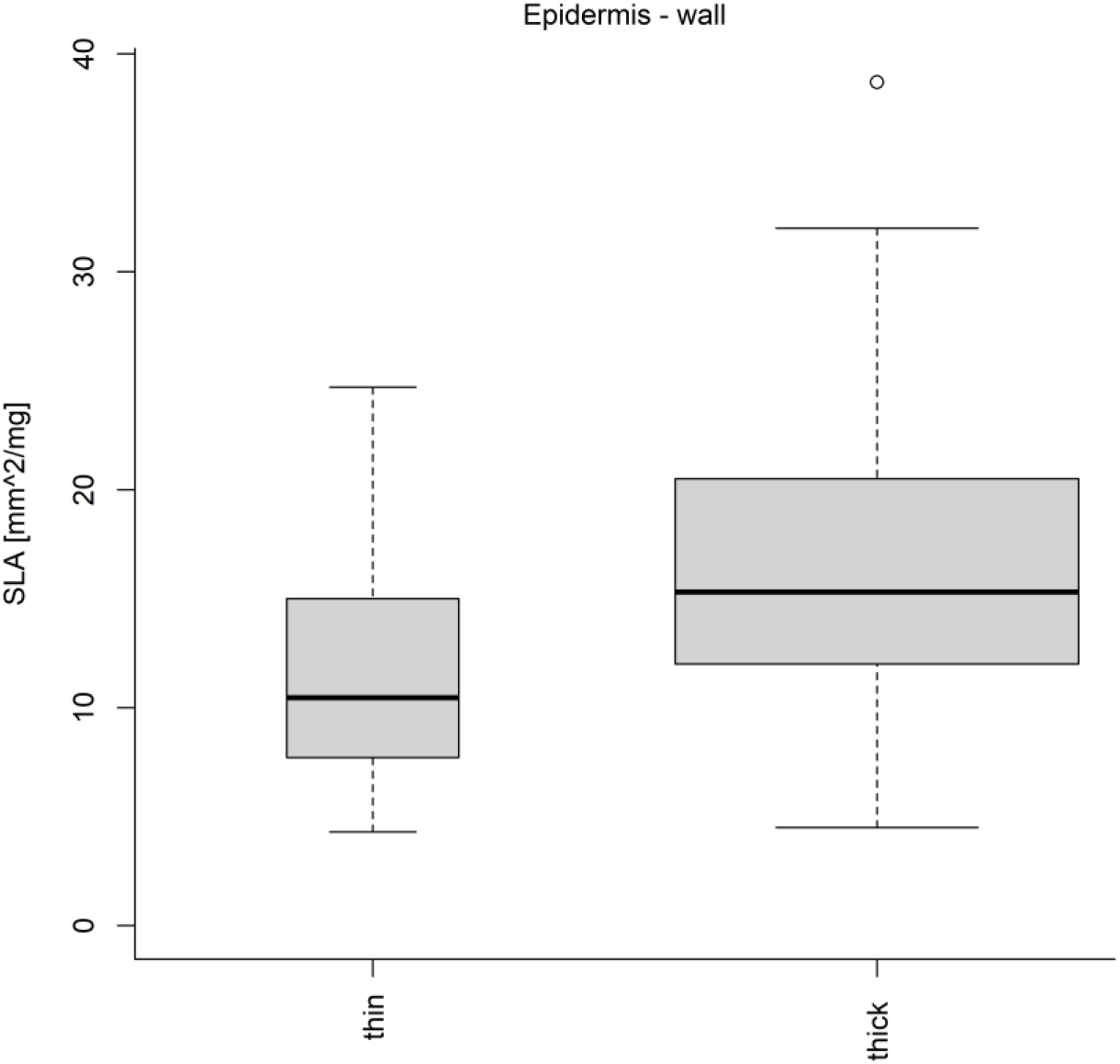
Relationship between SLA and type of epidermis wall. The width of boxplots corresponds to the square root of the number of observations in a particular group.

Leaf area accounted for the most variation in epidermal trichomes (Figure 2), which are either unicellular (42%), multicellular (19%), or absent (39%). Unicellular and mostly non-glandular trichomes predominated in small trees and shrubs with small LA and high leaf thickness (high LDMC) such as *Buxus sempervirens, Cydonia oblonga, Nerium oleander, Prunus* spp, and *Salix* spp (Figure 4). Petiole trichomes were missing in large trees with large and softer leaves such as *Fagus sylvatica, Aesculus hippocastanum*, and *Robinia pseudoacacia*. Less represented were branched non-glandular trichomes (*Hippophae rhamnoides, Olea europaea, Quercus coccifera, Rhododendron ferrugineum*), simple glandular trichomes (*Corylus avellana, Rhus typhina, Vitex agnus-castus*), and branched glandular trichomes (*Buddleja davidii*) (see also Supplementary Material Figures S6-S14).

The main mechanical support was achieved through the formation of collenchyma at the periphery of the petioles and stabilizing cortical fiber bands (Figures 3). Variation in collenchyma was mainly related to LA and temperature, with lamellar collenchyma found in 70% of species (see also Supplementary Material Figures S15-S21), mostly in shrubs and small trees from colder and wetter habitats with smaller leaves such as from Ericaceae, Cornaceae, Grossulariaceae, and Aquifoliaceae, while collenchyma tended to be missing in taller trees with large leaves such as *Populus tremula, Paulownia tomentosa*, and *Juglans regia* (Figure 4). Variation in cortical fiber bands was primarily related to plant height (Figure 3), with fiber bands continuous (36% of taxa), discontinuous (36%), or absent (28%). Fiber bands were largely absent in smaller trees and shrubs from Cornaceae, Aquifoliaceae, Rosaceae, Rhamnaceae, Grossulariaceae, and Oleaceae families, while continuous cortical fiber bands were massively developed in closely related large trees from Fagaceae (*Quercus, Fagus, Castanea*), Sapindaceae (*Acer*) and all Fabaceae (*Amorpha fruticosa, Ceratonia siliqua, Cercis siliquastrum, Gleditsia triacanthos, Glycyrrhiza glabra*, and *Robinia pseudoacacia*, Figure 4).

### 3.2. The arrangement of vascular bundles proportion and structure of the conductive system

The structure of vascular bundles was mainly related to temperature and SLA (Figures 3 and 8) and can be divided into three larger categories including i) crescent-shaped of phloem and xylem was found in 51% of studied taxa, mostly shrubs and smaller trees with though leaves (low SLA) such as *Laurus nobilis, Buxus sempervirens, Nerium oleander, Phillyrea angustifolia, Prunus mahaleb* and *Loiseleuria procumbens*; ii) phloem and xylem encircling vascular bundles were found in 41% of species, mostly large trees with softer leaves (*Quercus, Acer, Gleditsia, Juglans, Robinia*); and iii) multiple internal stems were found in (4%) of the species, *Populus* spp and *Salix aurita* (Figures S22-33). In vascular bundles, xylem predominated (76% of species) over phloem (15% of species) and sclerenchymatous tissue (9% of species). Phloem dominating over other tissue types was more frequently found in smaller trees (*Malus, Prunus, Rhus, Salix*) and shrubs (*Lonicera alpigena, Rhododendron ferrugineum*), while sclerenchyma predominates over xylem and phloem in larger trees from warmer and drier habitats unrelated groups such as in *Liriodendron tulipifera, Magnolia x soulangeana, Glycyrrhiza glabra* and *Quercus coccifera* (Supplementary Material Figures S34-S41). Xylem had distinct earlywood and latewood in 66% of studied taxa. Most of the studied species had lignified xylem with fibers composed of small cells (72%), mostly taller trees (Figure 4), while xylem without fibers tended to occur in smaller trees and shrubs across different families but all belonging to Rosopsida (Anacardiaceae, Araliaceae, Aquifoliaceae, Caprifoliaceae, Fabaceae, Rosaceae, Trochodendraceae) (Supplementary Material Figures S22-S33).

**Figure 8.**
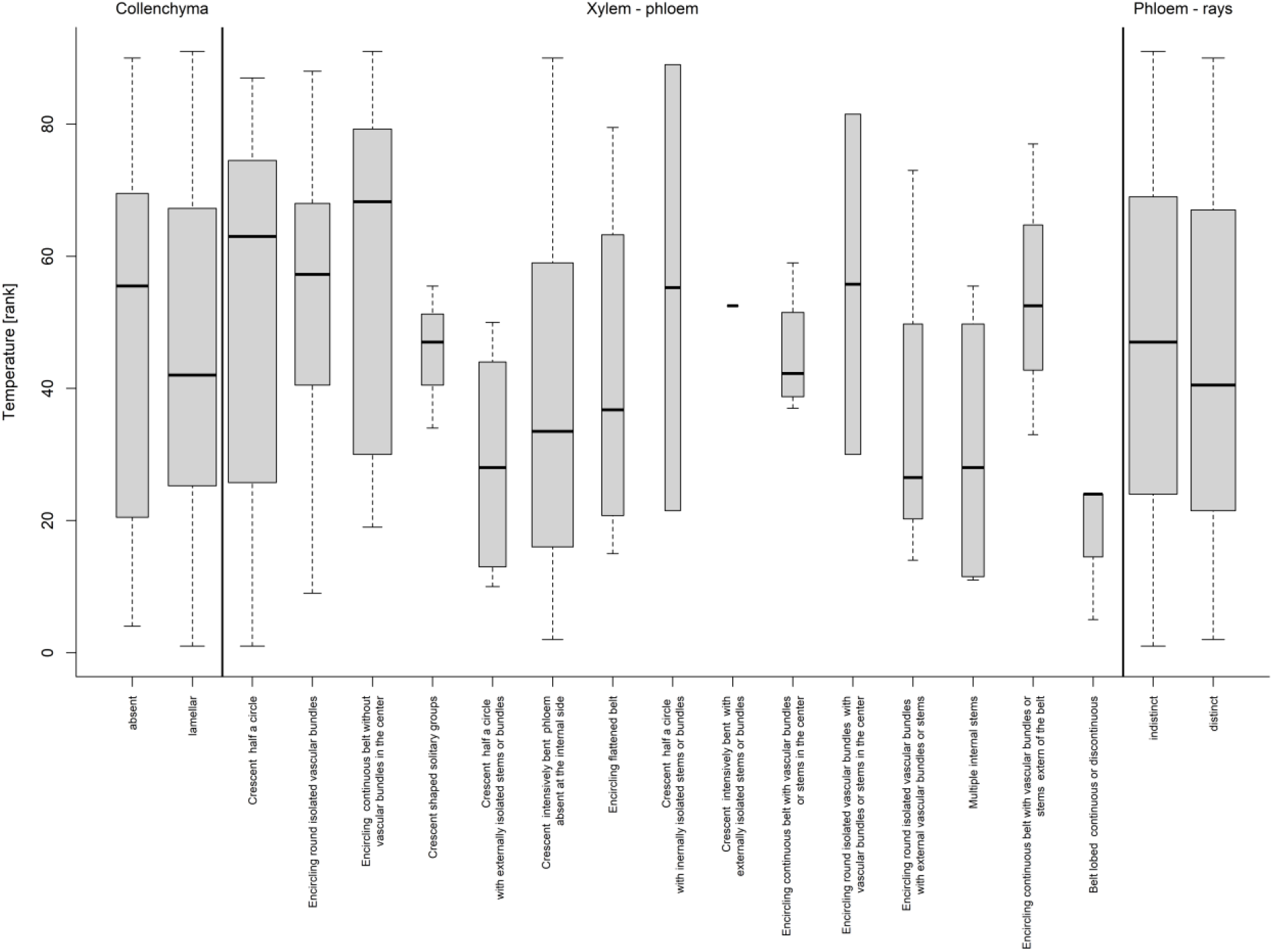
Relationship between temperature and selected anatomical traits. Width of boxplots corresponds to square root of number of observations in particular group

The arrangement of vessels in the xylem was best explained by variation in LA (Figures 2 and 3). The vessels are primarily radially distributed among studied species (71%) and less frequently irregularly distributed (29%) (Figure 2). Radially distributed vessels were mostly smaller than 20 µm in diameter and were typically found in species with smaller and thicker leaves such as *Nerium oleander, Phillyrea angustifolia, Buxus sempervirens, Arbutus unedo* (Supplementary Material Figures S22-S33), while irregularly distributed vessels were often larger than 20 µm in diameter and are found in higher trees with softer leaves as *Quercus* and *Populus* but also lianas such as *Clematis vitalba* and *Hedera helix* (Figures 2 and 4). Variation in vessel diameter was primarily related to leaf size variation (Figure 5). Narrow vessels (10-20 µm) occurred in 56% of taxa, mainly in shorter trees and shrubs with smaller leaves, while higher trees with bigger leaves such as *Quercus petraea, Juglans regia, Aesculus hippocastaneum* and lianas (*Parthenocissus tricuspidata* and *Vitis vinifera*) have wider vessels (30-50 µm, Figure 4). The narrowest vessels in petioles are typically found in small shrubs of cold and wet habitats such as *Rhododendron ferrugineum* and *Vaccinium vitis-idaea* or tree species from seasonally dry climates such as *Phillyrea angustifolia, Quercus coccifera, Ilex aquifolium, Gleditsia triacanthos*, and *Arbutus unedo* (Supplementary Material Figures S42-S51). Most of the studies species have phloem composed of small cells (72%), while the proportion of species with large phloem cells is the highest in small trees restricted to few families like Rosaceae (*Malus sylvestris, Mespilus germanica, Pyrus communis, Sorbus aria, Sorbus aucuparia*), Salicaceae, Sapindaceae, and Grossulariaceae. Phloem rays are distinct (49% of taxa) and indistinct (51%), influenced by temperature (Figure 8). The highest share of species with distinct phloem rays includes smaller trees from Betulaceae, Salicaceae, and Rosaceae families (Figure 4).

### 3.3. Crystal and Secretory elements

Variation in secretory elements and crystal was primarily related to plant height and precipitation, respectively (Figure 3). Distinct crystal types (prismatic, druses, acicular, sand) were found mainly in shorter taxa from drier habitats with smaller and thicker leaves (small SLA and high LDMC, Figure 3). The majority of species had both prismatic crystals and druses (46%), followed by petiole with only druses (26%), only prismatic crystals (9%), acicular crystals (6%), sand (4%), and druses and sand (2%). Acicular crystals are found in species from closely related families from order Lamiales (Lamiaceae, Oleaceae, Paulowniaceae) such as *Olea europaea, Syringa vulgaris, Paulownia tomentosa* and *Vitex agnus-castus*. Sand crystals are found in *Hippophae rhamnoides, Laurus nobilis, Phillyrea angustifolia* and *Sambucus* spp. Secretory elements were found only in 12% of the species. Secretory ducts occur in 7% of the species, including small trees (*Myrtus communis, Rhus typhina*), shrubs (*Cotinus coggygria, Frangula alnus, Pistacia lentiscus*), lianas (*Hedera helix*) and large trees (*Tilia platyphyllos*). Slime content is present in 5% of taxa, exclusively in trees such as *Betula pendula, Prunus avium* and *P. mahaleb* and *Ulmus glabra*.

## 4. Discussion

We presented here morphological and anatomical trends in petiole traits of woody species, as a first attempt to provide a broader view on the variation in petioles, which are a key organ supporting the main photosynthetic machinery in leaf blades and playing an essential role on the hydraulic pathway within the plant. Our evaluation improves our understanding of how variation in petiole morphoanatomical traits is driven by the plant height and leaf characteristics, as well as gradients of temperature and precipitation.

We found that longer petioles occurred in larger leaves (higher LA) from taller tree species that grow in warmer regions. In contrast, shorter petioles were present on smaller leaves (lower LA) from shorter trees and shrubs from drier regions. The general advantage of having longer petioles, often bigger than leaf blade area, is to provide radical changes in leaf orientation and thus optimized light-harvesting (Takenaka 1994; Niinemets et al. 2004). In other words, the variation of petiole length, combined with leaf blade dry mass, directly influences leaf angle, which may reduce the shading of basal foliage by adjacent leaves and thus increasing light-interception (Niklas 1991; Takenaka 1994; Niinemets and Fleck 2002; Niinemets et al. 2004). The relation between petiole length and plant height also raised questions associated with wind interference. Considering that longer petioles are more flexible to strong winds (Vogel 1989) and that the wind speed increase with height in the canopy (Niinemets and Fleck 2002), we would expect that longer petioles wiould be found in taller trees. Indeed, variation in petiole length is considered to be a strategy to cope with the vertical gradient of wind speed on trees, reinforcing the tendency we observed here.

Additionally to petiole length, petiole cross-sectional geometry (outline) is also an important trait that influences leaf response to wind stresses and leaf self-holding (Niklas 1991, 1996; Niinemets and Fleck 2002; Faisal et al. 2012; Louf et al 2018). In our observations, we identified five different petiole outlines, including circular, with indentation (terete petiole), flattened horizontally and vertically, and polygonal. We found that circular outlines tend to occur in longer petioles from taller trees, while petioles with an indentation tend to be more frequent in shorter petioles from smaller trees. This fact certainly represents an important morphological trait to understand trends in petiole biomechanics. However, because of the scarce information on this topic, it is difficult to evaluate in detail the way outlines influence leaf torsion and bending under wind exposure. Nevertheless, the previous report showed that petioles with a vertically flattened outline, typical of *Populus* species, promote the reduction of leaf torsion by the wind when compared to petioles with indentation (Niklas 1991). Our findings call for more experimental studies to better understand how petiole geometry influences leave flexibility and support among trees.

Despite morphological trends, the main adaptive responses of petiole occurred anatomically and were directly influenced by leaf area. In general, larger leaves would require petioles with appropriate anatomical qualities to provide more leaf support and flexibility to handle more intense mechanical stresses (Vincent 1982; Niklas 1990). In that case, supportive tissues like collenchyma, sclerenchyma, and xylem would provide these qualities. Indeed, we showed that collenchyma and sclerenchyma tend to be more frequent in longer petioles. Both tissue types are characterized by thick cell walls and are known to provide distinct mechanical properties. While collenchyma has soft cell walls, with viscoelastic properties that permit bending without compromising the internal structures, sclerenchyma has lignified and more rigid cell walls, which support and prevent damages to the adjacent tissues. We observed that collenchyma was located on the periphery of petioles, while sclerenchyma was associated with vascular bundles, which is in alignment with the typical distribution in petioles (Evert 2006). Moreover, distinct parts of the leaf (midrib, leaf border and leaf apex) are characterized to have both tissues more well developed to better hold mechanical stress rate (Niklas 1991; Evert 2006). In the swollen base of petioles, for instance, the presence of collenchyma is abundant and provide more plastic movements according to the wind (Niklas 1991; Evert 2006), while sclerenchyma is especially located next to the phloem, preventing the rupture of the thin cell walls due to mechanical and drought stress (Esau 2006).

Since leaf anatomy is extremely adaptable to environment conditions (Metcalf and Chalk 1950; Krober 2015; Stojnić et al. 2016) and intimately linked with temperature and water gradients (Bussotti et al. 1995; Gravano et al. 1999; Doria et al. 2019), the proportion of tissues may emphasize the conditions to which plants are exposed. However, anatomical variation is mainly described for leaf blades and less information is available for petioles. We observed that shorter petioles from drier environments have smaller epidermal cells and thicker cuticle, which is aligned with the features typically found for leaf blades exposed to similar conditions. The development of both traits is generally considered a strategy to minimize water loss via transpiration (Porsch 1926; Ennajeh et al. 2010; De Micco and Aronne 2012). Indeed, it is well known that the combination of these anatomical traits together with the reduction of leaf size represent xeromorphic features that are frequently observed in leaves from Mediterranean conditions, exposed to water deficiency, high temperatures and light intensity (Castro Diez et al. 1997; Rotondi et al. 2003). In contrast, our findings showed a distinct pattern for sclerenchyma in petioles. We observed that a higher proportion of sclerenchyma was present in longer petioles of taller trees that occurred in environments with higher temperatures but with not water deficiency. This fact indicates firstly that the strongest role of sclerenchyma in petioles is probably to provide mechanical support, and secondly, that petiole length has a stronger influence in sclerenchyma development.

Concerning xylem traits, our analysis showed that vessel diameter is also linked with leaf area, and again with petiole length. While vessels with bigger diameter are found in longer petioles (with larger leaves), narrower vessels are developed on shorter petioles (with smaller leaves). Indeed it is known that leaf vascular properties are strongly determined by leaf morphology but consistent responses are also connected with climatic variables (Sellin and Kupper 2007; Scoffoni et al. 2008, Sanginés de Cárcer et al. 2017, Kardošová et al. 2020). Even though we observed that xylem vessels tend to enlarge with higher levels of precipitation and temperature, the strongest relationship was still found with leaf size. This tendency is aligned with the associations between vessel widening and increasing leaf area previously made for *Fraxinus americana, Quercus robur, Acer pseudoplatanus* and other trees (Nicklas 1992; Coomes et al. 2008; Lechthaler et al. 2019; Levionoides et al. 2020). In these studies, the authors analyzed single species or small groups of plants, but here we show that the same trend occurred when a larger group of species is considered. It is known that longer petioles can only maintain larger leaves with wider vessels (Coomes et al. 2008; Levionoides et al. 2020) since the reduction of vessel diameter would represent diminished efficiency of water transport and thus reduction of gas exchange and carbon assimilation (Sack et al. 2003; Brodribb 2009; Jordan et al. 2013; Scoffoni et al. 2016; Levionoides et al. 2020). But it also suggests that the longer the leaf the wider its vessels at the middle of the petiole, the location from where the samples were collected for this study. This is in line with previous observations of vessels widening within the leaf, from the very narrow vessels at the end of the sap path close to the stomata towards the wider vessels at the base of the leaf (Rosell et al. 2017, Rosell et al. 2019).

Another interesting anatomical trend showed here is the presence of different cell types in vascular tissues, which may be considered relevant for the taxonomy of certain groups. Despite the high plasticity of leaf anatomy, different studies have proven that vascular patterns and superficial traits (epidermis, cuticle, trichomes) in petioles have an important value for the taxonomy of distinct groups (Hare 1944; Kocsis and Borhidi 2003; Noraini et al. 2016; Talip et al. 2017; Anu ad Dan 2020; Karaismailoğlu 2020). Our findings showed that in small trees and shrubs from some families (Anacardiaceae, Araliaceae, Aquifoliaceae, Caprifoliaceae, Fabaceae, Grossulariaceae, Rosaceae, Salicaceae, Sapindaceae, Trochodendraceae) the phloem has scattered bigger parenchymatic cells and the xylem is fiberless. Even though both traits have not been yet considered taxonomically relevant in petioles, in woody stems the patterns of parenchyma and the presence and absence of fibers are defined as the most conspicuous traits to characterize species (Chattaway 1953; Roth 1981; Archer and van Wyk 1993; den Outer 1993). Moreover, preview studies showed similar characteristics on the midrib of the same species (Săvulescu and Luchian 2009; El-Alfy et al. 2011; Koçyiğit et al. 2015; Jušković et al. 2017). Even though anatomical changes occur along the petiole and at the beginning of the midrib (Sack and Holbrook 2006), both traits seemed to have the potential for taxonomy. Nevertheless, further investigations are needed to evaluate their constancy and relevance for these species the study using "only” 95 woody species of the highly species-rich functional group of trees and shrubs and the geography constraints (Europe) excluding the tropics.

## 5. Conclusion

The structural traits of petioles we evaluated in this study included morphological and anatomical aspects that characterized their shape, stiffness and hydraulic potential. Our analysis was based on 95 major woody species from Europe representing one of the first attempts to find trends for the broad and diversified group of woody plants. Our results showed that leaf area has the strongest influence on petiole anatomical traits, more than temperature, precipitation, and plant height, which emphasize the supportive and mechanical role of this part of the leaf. Petioles tend to be longer and have a circular outline in larger leaves. Anatomically, mechanically supportive cells (collenchyma and sclerenchyma) tend to be more predominant and xylem vessels tend to have a bigger diameter in bigger leaves. These traits are aligned with our expectations, as bigger leaves represent more weight for self-holding and demand a more efficient vascular tissue. In the case of smaller leaves, petioles tend to be shorter, to have an outline with indentation and narrower vessels. Indeed, environmental factors may directly influence distinct plant organs, however, our results appear to reveal a different pattern for petioles that relates to the leaf itself. Since this study is constrained geographically to Europe and hence based on a limited set of woody species selected from otherwise highly species-rich functional groups of trees and shrubs, further investigations are needed to better understand the evolution of petiole structures in temperate and especially tropical woody species.

## Acknowledgments

J.D., A.L.F. and J.A. were supported by the Czech Science Foundation (Project: 21-26883S) and MSMT LTAUSA18007. AC was supported by a generous donation from Fritz H. and Elisabeth Schweingruber.

