## Supplementary Material for "Comparative anatomy of leaf petioles in temperate trees and shrubs"

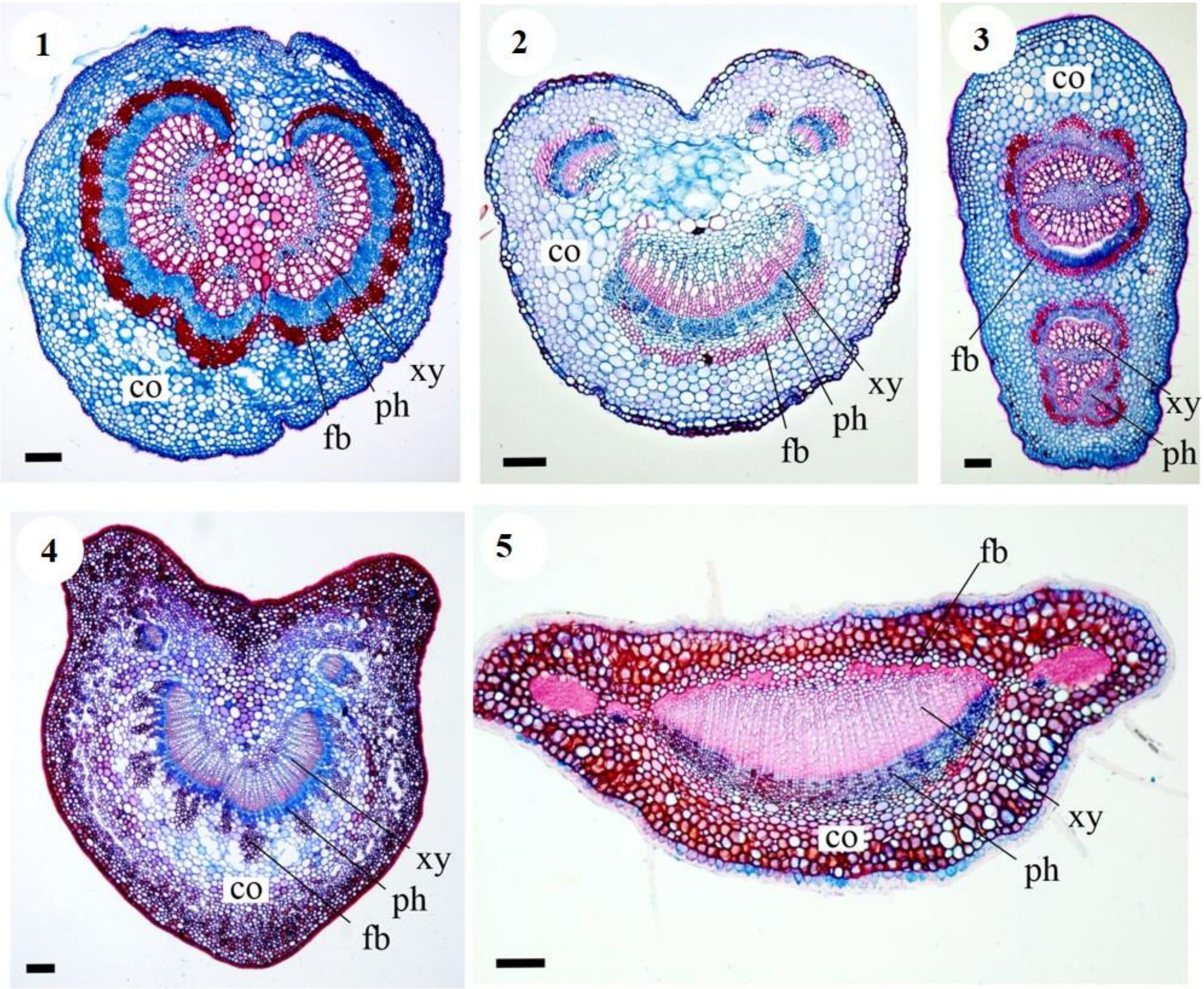


**Figures S1-S5.** Cross-sectional geometry (outline). 1. Circular in *Alnus cordata*. 2. Almost circular, with an indentation in *Coronilla emerus*. 3. Flattened vertically in *Populus tremula*. 4. Polygonal in *Prunus laurocerasus*. 5. Flattened horizontally in *Buxus semprevirens*. co = cortex; fb = fiber band; ph = phloem; xy = xylem. Scale bars = 200 µm.

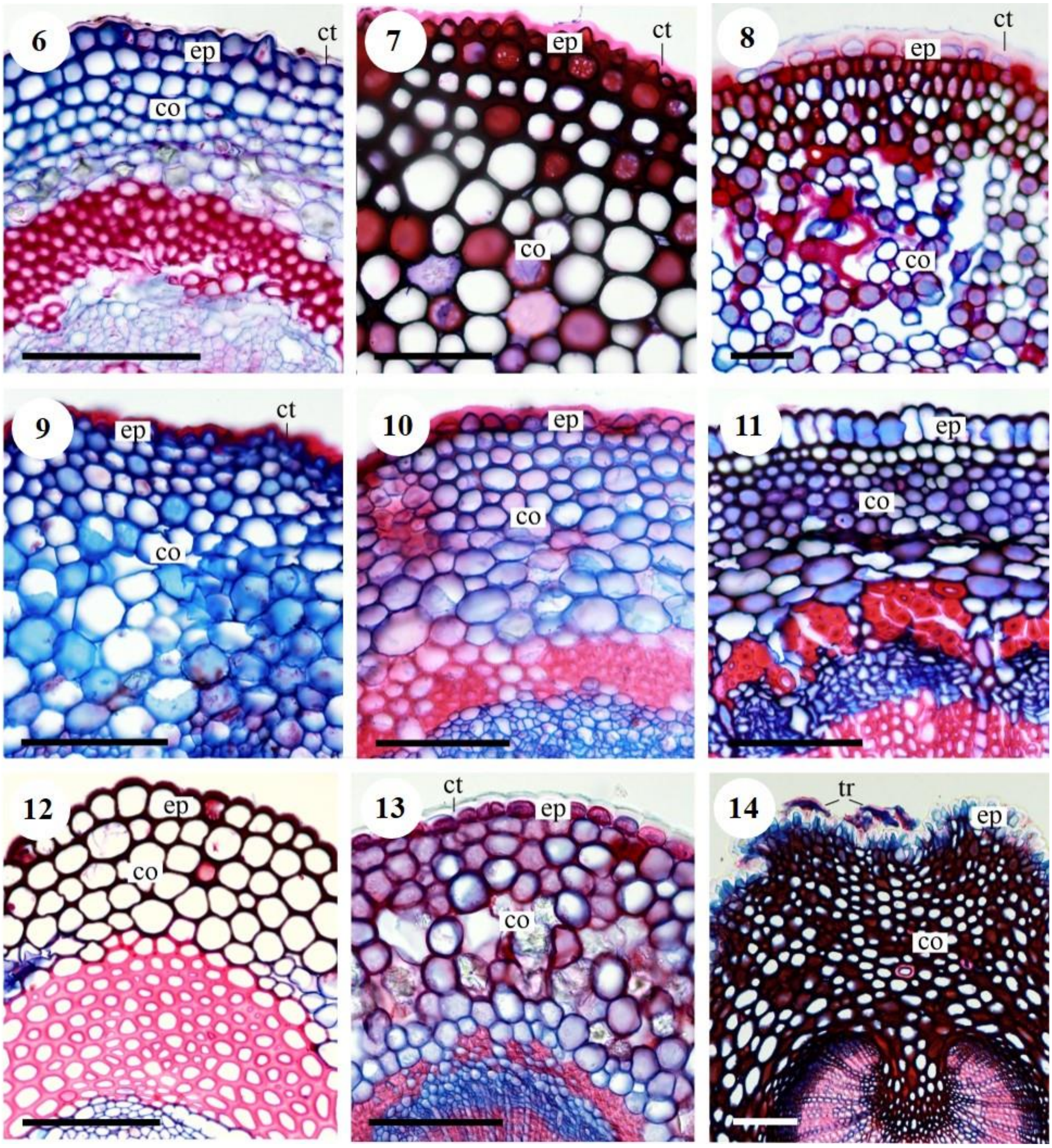


**Figures S6-S14.** S6-S8. Cuticule thickness: <1 µm in *Acer platanoides* (6); 2-5 µm in *Viburnum tinus* (7); > 6 µm in *Trochodendron araloides* (7). 9-11. Size of epidermal cells compared to cortical cells: smaller than cortical cells in *Salix alba* (9); similar to cortical cells in *Rosa arvensis* (10); bigger than cortical cells in *Capparis spinosa* (11). 12-13. Thickness of epidermal walls: thick in *Clematis vitalba* (12); thin in *Vaccinium vitis* (13). 14. Presence of trichomes in *Olea europaea*. ct = cuticule; ep = epidermis; co = cortex; tr = trichome. Scale bars = 100 µm.

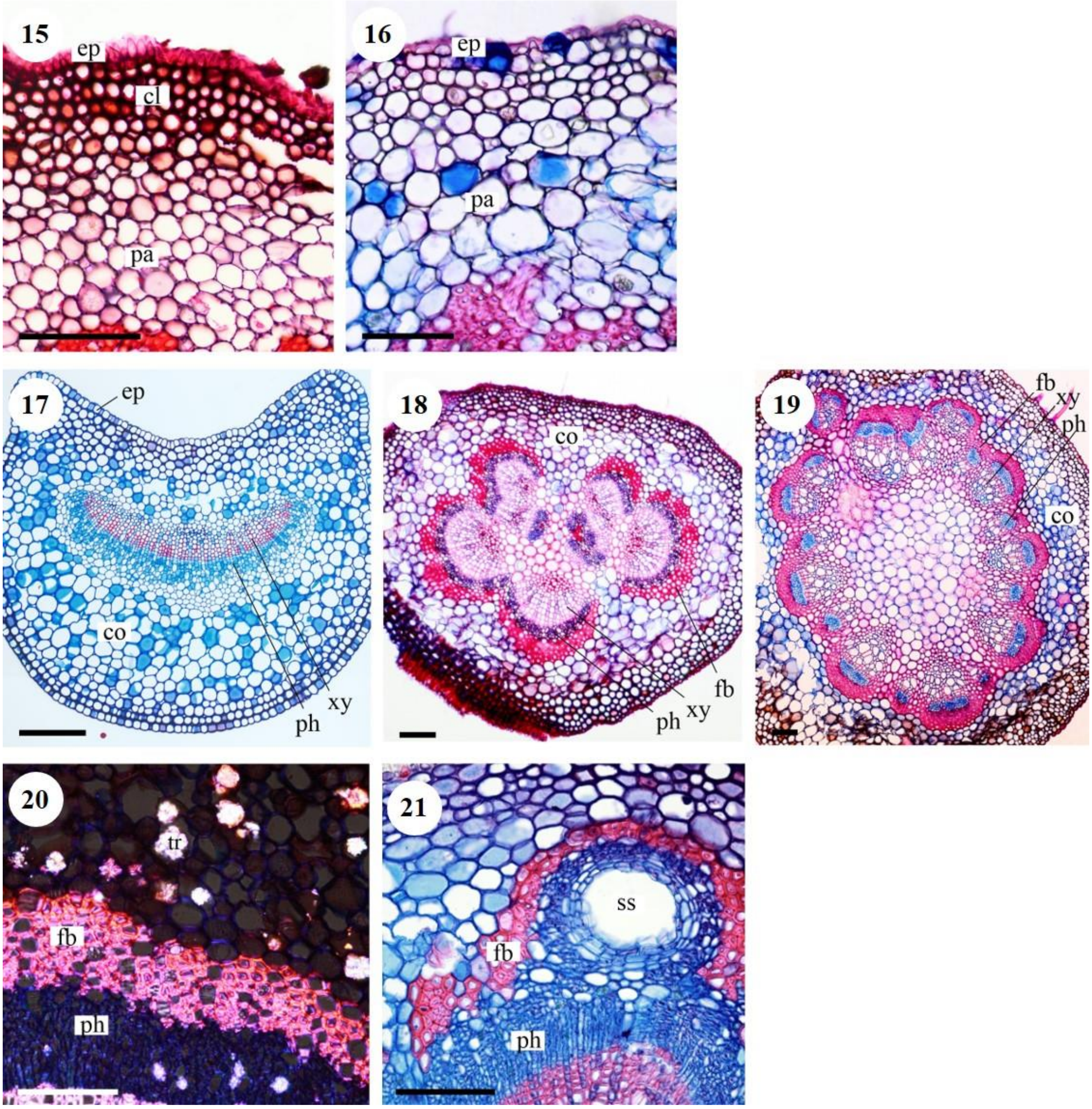


**Figures S15-S21.** Presence of collenchyma in *Malus sylvestris* (15). Absence of collenchyma in *Pyrus communis* (16). 17-19. Fiber band: absent in *Lonicera alpigena* (17); discontinuos in *Carpirus betulus* (18); continuous in *Magnolia* × *soulangana* (19). Presence of crystals in *Alnus alnobetula*. Presence of secretory structures in *Pistacia lentiscus*. ep = epidermis; co = cortex; cl = colenchyma; pa = parenchyma; fb = fiber band; ph = phloem; xy = xylem; cr = crystal; ss = secretory structure. Scale bars = 100 µm (15, 16, 20, 21), 200 µm (17-19).

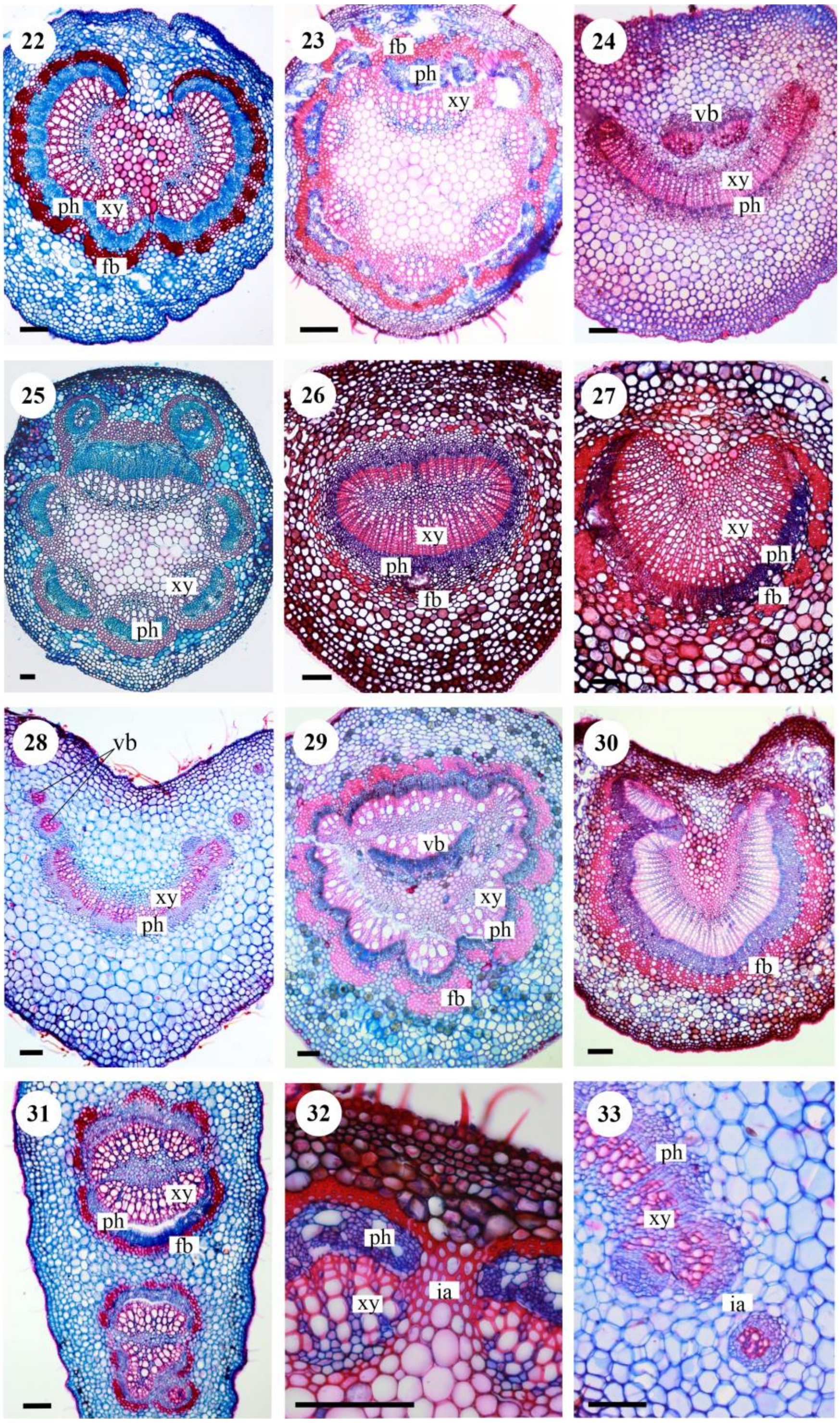


**Figures S22-S33.** S22-S31. Arrangement vascular system (phloem-xylem): lobed belt in *Alnus cordata* (22); encircling round isolated vascular bundles in *Acer campestre* (23); crescent half a circle with externally isolated bundles in *Cornus sanguinea* (24); encircling round isolated vascular bundles with external vascular bundles in *Acer platanoides* (25); encircling flattened belt in *Arbutus unedo* (26); crescent in half a circle in *Amelanchier canadensis* (27); crescent shaped with solitary groups in *Buddleya davidii* (28); encircling continuous belt with vascular bundles in the center in *Quercus rubra* (29); crescent shape and intensively bent in *Alnus alnobetula* (30); multiple internal rings in *Populus tremula* (31). The interfascicular area is lignified (32) or parenchymatic (33). vb = vascular bundles; fb = fiber band; ph = phloem; xy = xylem; ia = interfascicular area. Scale bars = 100 µm (32, 33), 200 µm (22-31).

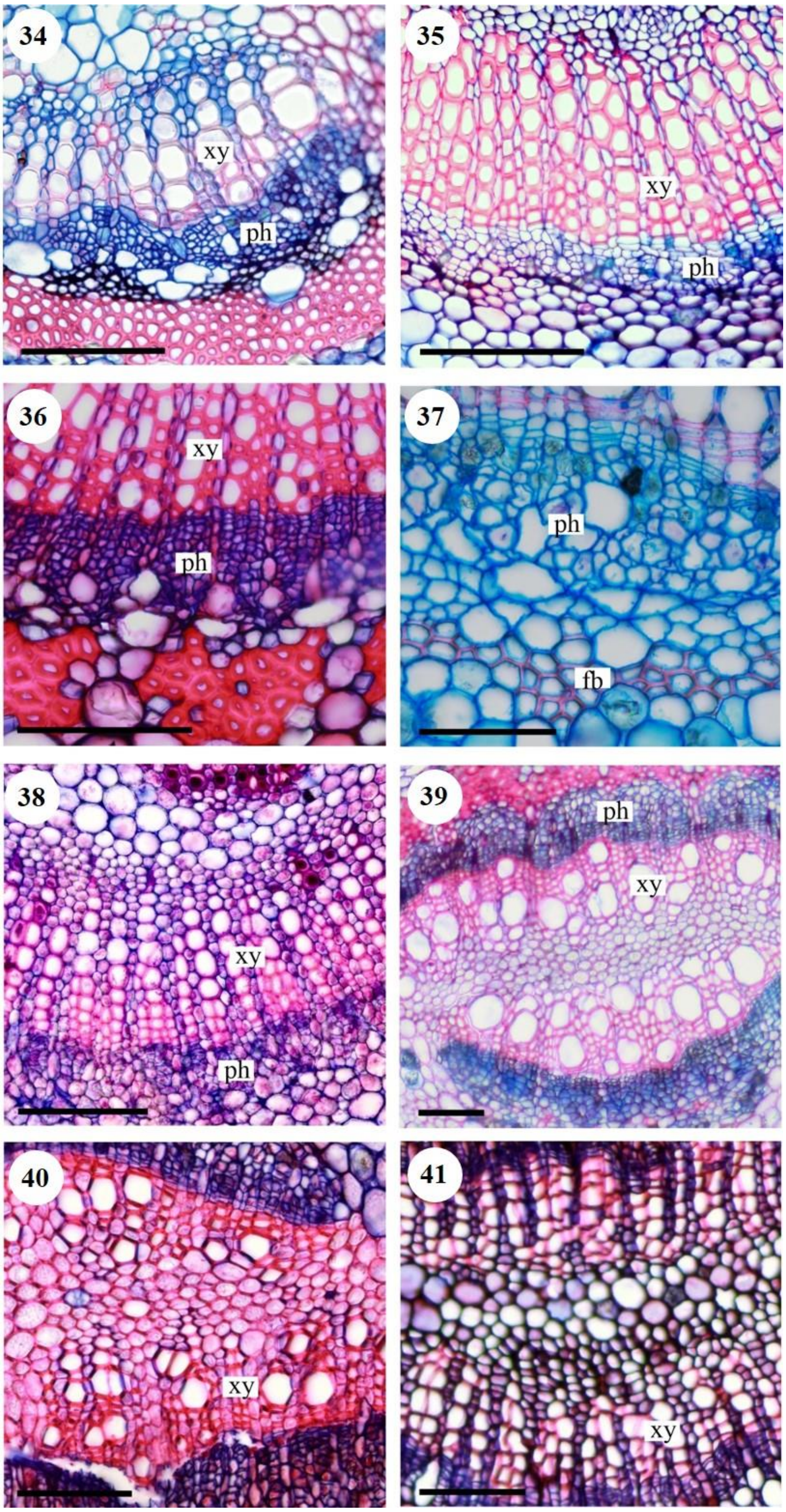


**Figures S34-S41.** Phloem have large cells in *Amorpha fruticosa* (34) and is without large cells in *Nerium oleander* (35); with visible rays in *Amelanchier canadensis* (36) and without rays in *Parthenocissus tricuspidata* (37). Xylem is fibersless in *Cornus sanguinea* (38) and with fibers in *Quercus rubra* (39). The arrangement of xylem vessels are irregular in *Quercus robur* (40) and radial in *Salix caprea* (41). ph = phloem; xy = xylem. Scale bars = 100 μm.

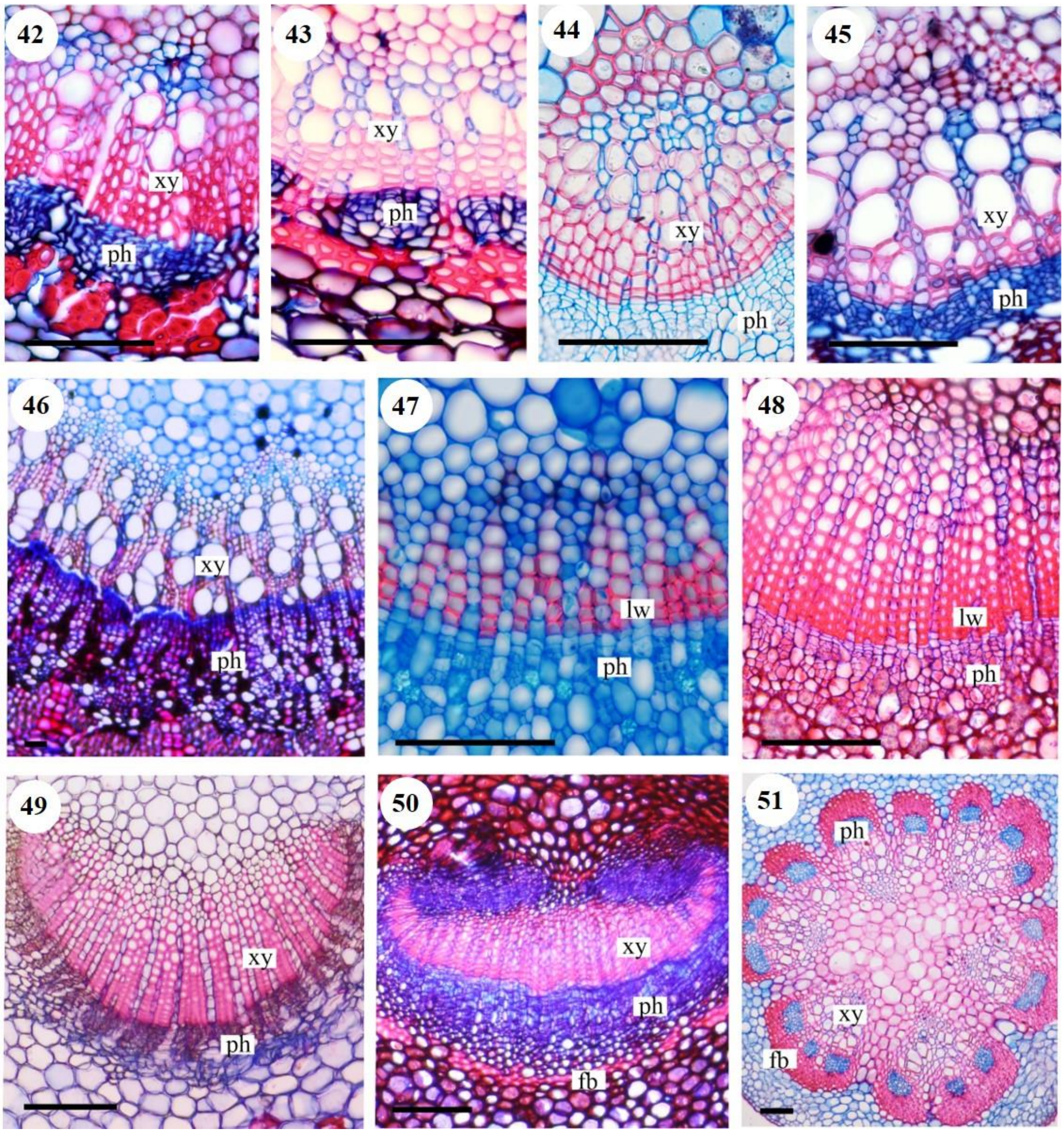


**Figures S42-S51.** Vessel diameter is <10 µm in *Hedera helix* (42), between 10-20 µm in *Acer pseudoplatanus* (43), between 20-30 µm in *Fraxinus excelsior* (44) and between 30-50 µm in *Capparis spinosa* (45). Latewood is indistinct in *Juglans regia* (46), distinct in *Lonicera alpigena* (47) and very distinct in *Cornus mas* (48). The predominant tissue is xylem in *Lingustrum vulgare* (49), phloem in *Myrtus communis* (50) and schlerenchyma in *Liriodendron tulipifera* (51). ph = phloem; xy = xylem. Scale bars = 100 µm (42-46), 200 µm (47-51).

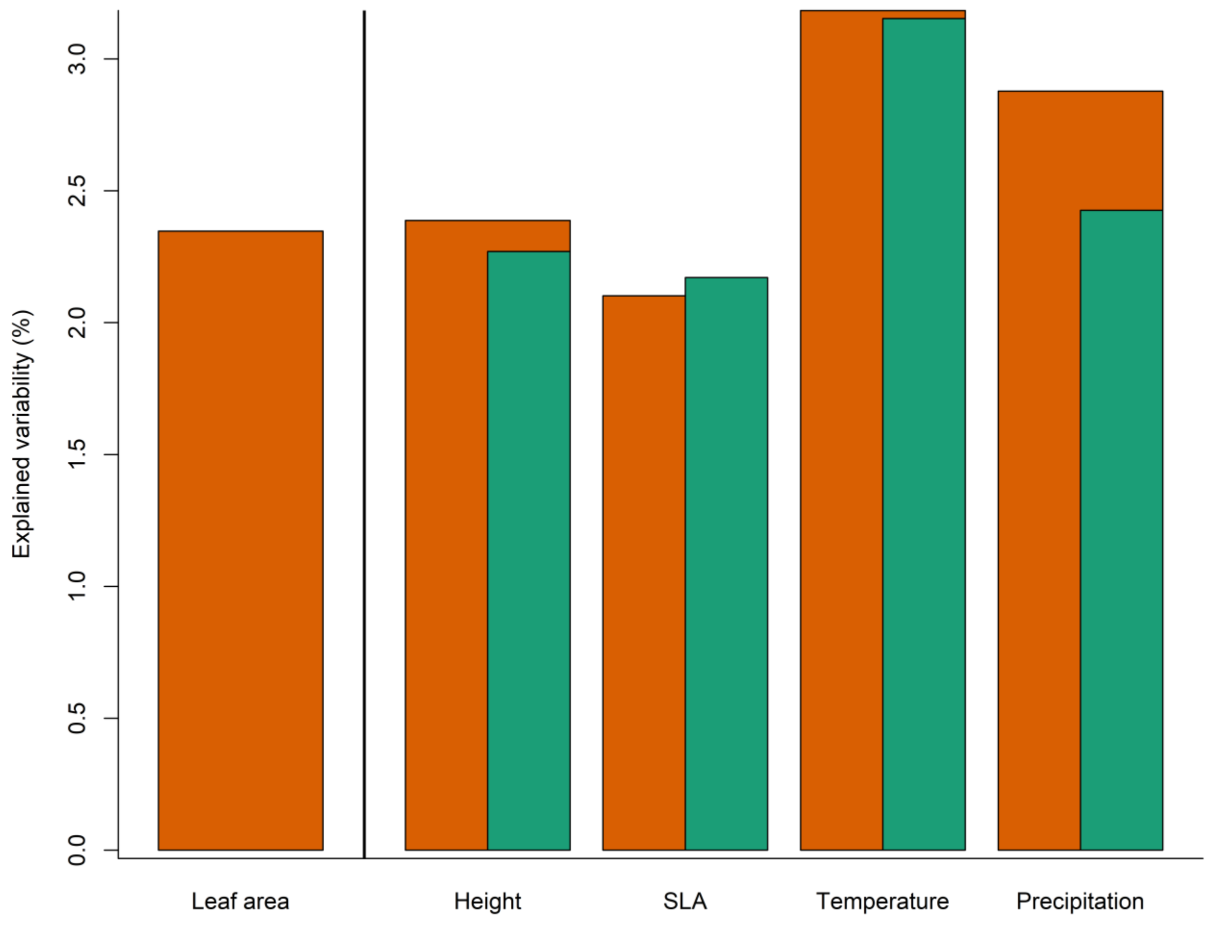


**Figure S52.** Explained variability of petiole anatomy by LA, height, SLA, precipitation, and temperature. Brown columns show explained variability accounting only for LA; green columns show explained variability after accounting for all the other predictors (including LA). LA = leaf area, SLA = specific leaf area.
